# Heritable cognitive and psychopathology factors in youth are predicted by brain fronto-temporal white matter pathway

**DOI:** 10.1101/172080

**Authors:** Dag Alnæs, Tobias Kaufmann, Nhat Trung Doan, Aldo Córdova-Palomera, Yunpeng Wang, Francesco Bettella, Torgeir Moberget, Ole A. Andreassen, Lars T. Westlye

## Abstract

A healthy transition from adolescence to adulthood relies on a continuous individual adaptation to a dynamic environment. Here, we employed data driven multivariate approaches to derive both general cognitive and psychopathology factors as well as brain phenotypes in children and adolescents in the publicly available PNC sample. We identified a distinct brain white matter pattern which proved central for prediction of heritable cognition and psychopathology scores, highlighting the importance of fronto-temporal connections for intellectual and mental development.

## Introduction

Adolescence is a highly sensitive period and a major window for adaptation and opportunities. Modulated by the temporal evolution of a range of gene-environment interactions, it also coincides with the emergence of many mental disorders^1^, which may reflect a costs of the potential for brain plasticity offered during this period. Several lines of evidence, including (1) substantial shared genetic contribution to mental disorders, (2) high rate of psychiatric comorbidity, (3) reports of a common underlying factor across diagnostic categories, on which high-scoring individuals are more likely to have cognitive impairments as well as compromised early-life brain function, all highlight the importance of identifying brain based intermediate phenotypes that may be shared across disorders and present before the onset of disease^1,2,3^. Crucially, the myelinated axons of the brain white matter (WM) form the structural backbone that enables functional integration and adaptation of neural networks, together termed brain connectivity. There is a long-held notion of brain dysconnectivity as a transdiagnostic model for psychopathology, where aberrant brain biology and synchronization modulate the risk for mental health issues^4^. Recent demonstration of a link between a specific pattern of cross-talk between brain networks and a general positive-negative mode of population variability across a range of behavioral, demographic and lifestyle measures, supports that the level of individual adjustment can be conceptualized along a single dimension with a common brain basis^5^. Also, another recent study support the existence of a general and heritable psychopathology factor in children and youth^6^. Here we replicate and extend those findings in the Philadelphia Neurodevelopmental Cohort (PNC^7^), by showing that a general cognitive and a general psychopathology factor are heritable with a shared genetic contribution, and are associated with a distinct pattern of white matter microstructure and connectivity in the developing human brain.

The microstructural and connectivity properties of the human brain WM undergo profound changes throughout childhood and adolescence, with a protracted maturation continuing well into the third decade^8, 9^, and thus play a paramount role in the development of a healthy mind. Whereas the boundaries of this maturational potential are likely regulated by genetic predispositions, experiences and environmental perturbations mediate the individual development of cognitive and mental characteristics^10^. The intricate biological processes shaping the brain connectome during neurodevelopment is highly coordinated across the brain, yet WM development in childhood and adolescence also display anatomically differentiated trajectories, suggesting regional variation in rates and timing of maturational changes^8^. Hence, when considered from a system-level perspective different brain circuits may compete for common resources, and the cognitive and clinical correlates may be modulated by the relative balance between different networks subserving distinct functions. Indeed, diffusion weighted imaging (DWI)-based indices of brain tissue organization may reflect the relative balance between differentially affected crossing fiber populations^11^. Consequently, fusion of multimodal brain imaging indices, which allows for a joint consideration of different brain characteristics, may yield more biologically interpretable patterns of brain microstructure and connectivity sensitive to the development of cognitive and mental health.

Here, and in line with the aims of the NIMH Research Domain Criteria Project (RDoc) for mental disorders^12^, we targeted a data driven multivariate delineation of both brain and behavioral phenotypes. Scores from a diverse battery of cognitive tests and clinical questionnaires for 6,487 individuals in the PNC were decomposed to form and isolate both general and specific phenotypic traits, including general cognitive (gF) and general psychopathology (mean-ICA) factors. In addition, we investigated seven independent clinical components each reflecting empirical clustering of specific psychopathology symptoms (Supplementary Fig. 1). We used Genome-wide Complex Trait Analysis (GCTA)^13^ to estimate the lower-bound heritability of the general cognitive and general psychopathology phenotypes based on variation in common single-nucleotide polymorphisms (SNP-heritability) in a subsample of 2963 individuals of European descent. We also estimated the genetic correlation between the traits and its precision by means of resampling. In a subset with available diffusion-weighted MRI data (n=813), we used linked ICA (LICA)^14^ to decompose eight WM maps per individual reflecting different microstructural and connectivity properties (fractional anisotropy [FA], axial diffusivity [L1], radial diffusivity [RD], mean diffusivity [MD], mode of anisotropy [MO], dominant fiber direction [f1], nondominant fiber direction [f2] and probabilistic tractography density [CD] maps) into 20 independent components. Each component is characterized by group level spatial maps (one per WM index), and corresponding subject weights (one per participant) reflecting individual participant’s contribution to the observed brain patterns. These weights were used in multivariate cross-validated prediction analysis as well as to test for univariate associations with age, cognition and psychopathology.

## Methods

### Sample and exclusion criteria

The analysis was based on the publicly available Philadelphia Neurodevelopmental Cohort^7, 15^ (PNC, access obtained with permission #8642, project title: Neurodevelopmental brain networks: Integrating multimodal imaging, cognition and genetics). The institutional review boards of the University of Pennsylvania and the Children’s Hospital of Philadelphia approved all study procedures, and written informed consent was obtained from all participants. We accessed diffusion weighted imaging (DWI) data from 883 individuals. After removal of all participants with severe or major medical health conditions (n=44, rating of medical condition was performed by trained personnel in the PNC study team), and datasets that did not pass quality assessment (n=91), the final MR-sample comprised 748 individuals (343 males) aged 8.7 to 22.6 years (mean: 15.1 years, SD: 3.3 years). We accessed cognitive and clinical data for 6,487 participants for decomposition of neuropsychological and clinical scores using PCA and ICA. Of the 748 participants included in the MR-sample, 729 had available clinical data. We accessed genetic data for 4,450 participants of European descent.

### MRI acquisition

DWI scans were acquired at University of Pennsylvania^15^ on a 3T Siemens TIM Trio scanner, acquired using a twice-refocused spin-echo (TRSE) single-shot EPI sequence (FOV: 240 × 240 mm; matrix: 128 × 128 × 70; 64 diffusion-weighted directions; b = 1000 s/mm^2^; voxel resolution 1.875 × 1.875 × 2 mm).

### MRI Quality Assessment

We computed temporal-signal-to-noise ratio (tSNR) for all DWI datasets using scripts and cut-off values previously described for the PNC sample^16^. Out of the 839 datasets remaining after exclusion based on medical conditions, 91 individuals (10.8%) were excluded based on a tSNR cut-off of 5.7.

### Image processing

DWI data were processed using FMRIB's Diffusion Toolbox (FDT), part of FSL 5.0^17^, and included correction for motion and eddy current distortions using eddy^18^. FA, eigenvector and –value maps were computed by fitting a tensor model to the raw diffusion data and used to derive mean diffusivity (MD), radial diffusivity (RD) and mode of anisotropy (MO)-maps.

### Cross fiber modeling

Estimation of the probability distribution of the diffusion directions was performed with the GPU-accelerated version of bedpostx^19^, with up to three fibers estimated per voxel.

### Probabilistic fiber tracking

Whole brain probabilistic fiber tracking was performed using probtrackx2^20^. In line with a recent application in aging and dementia^21^, for each participant and from each voxel inside the native space whole brain seed mask, 100 pathways were sampled, resulting in a 3D-volume tractography-map per participant which were normalized by dividing with the total number of streamlines processed. Thus, the value in each voxel represents the likelihood that any streamline will pass through that voxel (connection density). We also performed a second probabilistic tracking analysis, sampling 5000 streamlines pr. voxel inside four ROIs in the insular region bilaterally, based on the results from the LICA.

### Spatial normalization

All maps were nonlinearly transformed to the FMRIB58_FA template and skeletonized using the FDT TBSS pipeline^22^. For visualization of skeleton-based results we used tbss_fill which thickens the skeleton for easier visual interpretation. Fiber orientation maps were warped and skeletonized using tbss_x^23^.

### Linked independent component analysis (LICA)

We performed data-driven decomposition of imaging features into independent components using FMRIB’s LICA (FLICA, http://fsl.fmrib.ox.ac.uk/fsl/fslwiki/FLICA), which models the intersubject variability across modalities^14, 24^. The model order was heuristically chosen at 20 components based on inspection of the spatial maps.

### Neurocognitive test battery and gF-score

All PNC participants completed a computerized neurocognitive battery^25^. We included performance scores from 12 neurocognitive tests (Supplementary Table 2) spanning executive control and WM, episodic memory, verbal and non-verbal reasoning, and social cognition in a PCA. All scores were adjusted for age before entering the PCA, and the first factor was extracted as a general measure of neurocognitive performance (gF).

### Clinical measures and mean-ICA score

A computerized protocol was used to assess symptoms of anxiety, mood, behavioral, eating and psychosis spectrum disorders with collateral informants for individuals below 18 years of age^25^. We included 129 clinical symptom score items covering 18 psychological clinical domains (Supplementary Table 3) for all the available participants in the PNC sample (n=6,487), excluding all follow-up and conditional items, and computed summary scores for each domain. 1,627 participants had missing values on one or several clinical items. For all but two participants which had missing values on all 129 clinical items, the missing values was replaced with the nearest-neighbor value based on Euclidean distance. The percentage of missing values for the 129 items ranged from 0−7%.

All available clinical item scores were submitted to independent component analysis (ICA) using ICASSO^26^, decomposing them into seven independent components. The model order of the ICA was chosen after testing several different model orders ranging from 3-15, each of which 100 permutations was run, and based on the observed independence and reliability of the resulting components. The mean subject weight across components were computed (mean-ICA) and used as a general measure of psychopathology. For comparison, we submitted the clinical variables to a PCA and extracted the first principal component (pF). The correlation between mean-ICA and pF was r=.97.

### Statistical analysis

Prediction of Age, gF and mean-ICA using LICA subject-weights was performed using shrinkage linear regression, with automatic estimation of the correlation shrinkage intensity^27^. We performed 1000 iterations each with a separate 10-fold cross-validation, before calculating the mean correlation across all iterations. Feature importance was assessed by correlation-adjusted marginal corelation (CAR)-scores^28^. To assess significance, for each trait we performed 10,000 permutations of the 10-fold cross validated prediction while randomly permuting the trait-scores. Univariate associations between LICA subject weights and cognitive and clinical variables were performed using linear models, and included age, gender and tSNR as covariates, and corrected for multiple comparisons using false discovery rate (FDR). All reported p-values are two-tailed unless otherwise specified. For visualization of age-curves of LICA subject weights for high and low scoring individuals (gF and clinical ICA-score) we used a cut-off of +/− 1 s.d. to form the following groups: gF high (n=114), gF low (n=75), mean-ICA high (n=76), mean-ICA low (n=115), IC2 high (n=126), IC3 high (n=80) and IC4-high (n=108). We then repeatedly fitted a smooth function with automatic estimation of the smoothness parameter^29^ and computed the mean and s.d. across 10.000 bootstraps for each of the groups.

### Genomwide complex trait analysis

Genotyping of the PNC sample^25^ was performed by the Center for Applied Genomics, at the Children’s Hospital of Philadelphia. We included data from the following platforms: Illumina OmniExpress (*n* = 1,241), Illumina Human-610 Quad (*n* = 1,895), Illumina HumanHap-550-v1 (*n* = 317), Illumina HumanHap-550-v3 (*n* = 997). The chip-genotyped SNPs overlapping across the platforms were extracted and used to form a data set to apply the following filters on: individuals with relatedness > .05 (n=296); removal of SNPs in the major histocompatibility complex (25:35 Mb region on chr6) and the inversion region of chr8 (7:13 Mb); SNPs with genotyping rate <99%, minor allele frequency < 5% and Hardy-Weinberg Equilibrium test p-value < .0000001; pairwise pruning of SNPs in linkage disequilibrium (r^2^ > 0.2, window of 5,000, step of 500). We used GCTA^13^ to create a genetic relationship matrix based on the remaining SNPs (n=49,774), before running univariate and bivariate REML-analysis to estimate SNP-heritability for gF and mean-ICA, as well as the genetic correlation between these traits, in participants with overlapping phenotypic and genetic data (n=2,946). Age, gender and platform were included as covariates. The genetic correlation (rG) between gF and mean-ICA-scores was estimated through sub-sampling by drawing 6000 random sub-samples (n=2,746) from the full sample (n=2,945) without replacement, before estimating the mean and standard deviation of rG across the sub-samples.

## Results

Common SNPs explained 18% (p<.05) of the individual differences in general cognition and 16% (p<.05) of the differences in general psychopathology (Supplementary Table 1). These two traits also showed a negative genetic correlation (resampling mean r=−.74; s.e.=.15, 99% CI [−1, −.36]; Supplementary Fig. 2). The LICA components captured distinct patterns of neurodevelopment (Fig. 1A). All components combined predicted age with high accuracy (correlation between observed and predicted age: r=.77, p<.0001; p-value obtained from non-parametric permutation testing, Supplementary Fig. 3). This confirms the high sensitivity of brain features for age-prediction in the PNC sample^30^, and was primarily driven by the first component, capturing spatially global variance in FA, RD and MD (LICA-01, Supplementary Fig. 4). Strikingly, the components also allowed for significant prediction of general cognition (r=.39, p<.0001) and general psychopathology (r=.24, p=.0001). These were both strongly driven by the same component (LICA-09) of which anatomical distribution and multimodal contribution is consistent with degree of crossing fibers in fronto-temporal connections (Fig. 1B).

**Figure 1.**
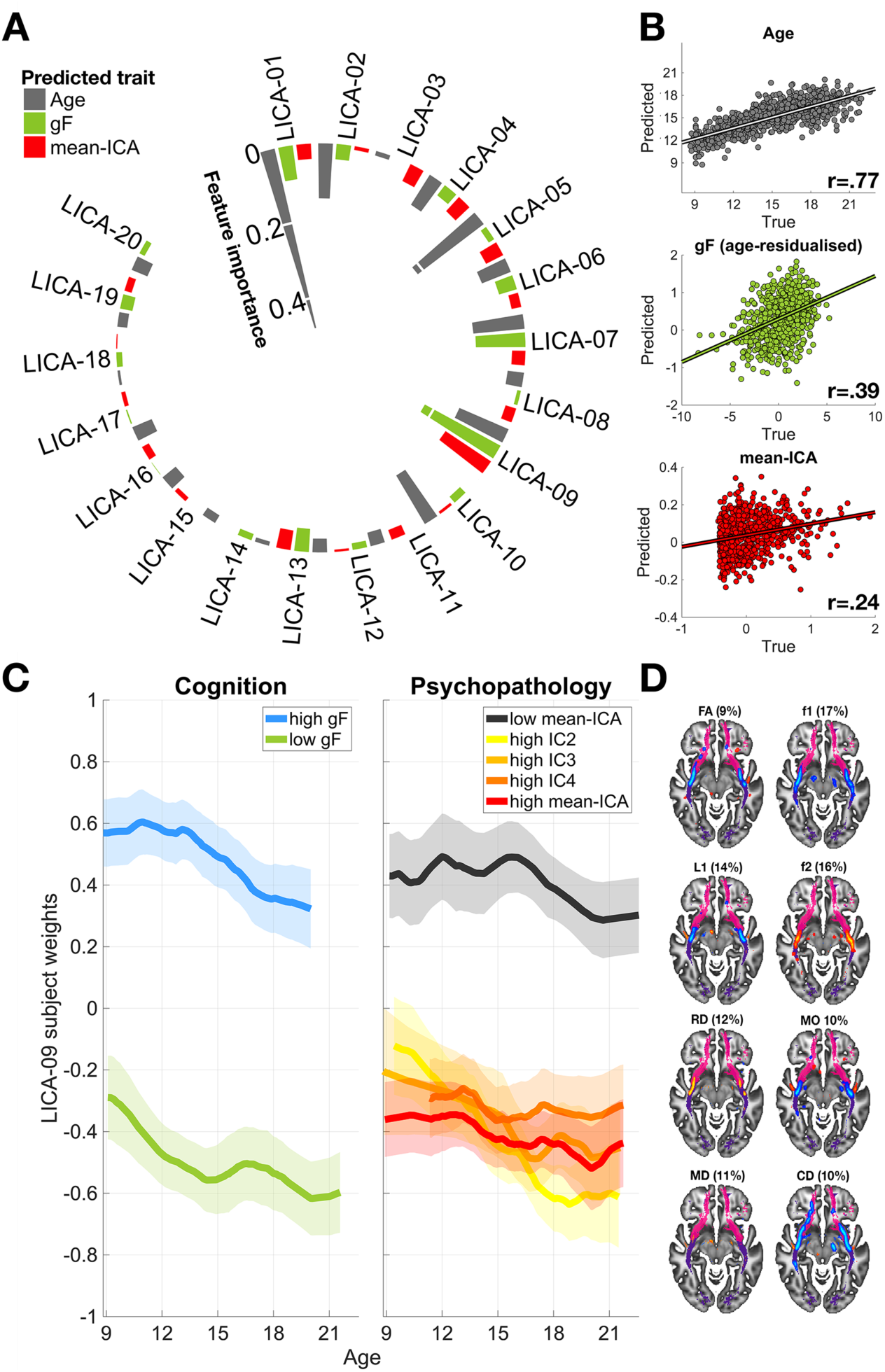
LICA brain components robustly predicted age, cognition and psychopathology **(A)** The circular plot shows the feature importance for each of the 20 components in cross-validated prediction of age, cognition (gF) and psychopathology (mean-ICA). While LICA-01 is the most important feature for prediction of age, LICA-09 is most important for prediction of cognition and mental health. **(B)** The scatter plots show the correlations between the true and predicted scores, all significant at p<.001 (FWE, 10.000 permutations). **(C)** The LICA-09 subject weights plotted as a function of age, and stratified by gF- and clinical ICA-scores. High- and low-scoring groups where defined as those scoring at least +/− 1 s.d. from the group average. Clinical IC2 represents high anxiety, while clinical IC3 and IC4 represent norm violating behavior and psychosis prodrome, respectively. Curves and shaded area represent the mean and s.d. of smooth function fitting across 10.000 bootstraps. **(D)** Brain maps represent the spatial weights for the diffusion MRI modalities of LICA-09, with percentages representing the relative modality contributions. Warm and cool colors represent positive and negative LICA-weights, respectively (z-threshold of +/− 3), overlaid on the UF (pink) and IFOF (purple) from the JHU WM-tractography atlas.

Specifically, the spatial foci of LICA-09 encompassed the intersection between the uncinate fasciculus (UF) and the inferior occipito-frontal fasciculus (IFOF)^31, 32^, with negative spatial weights for FA, f1, MO and CD and positive weights for f2 and RD. While an increase in FA is often interpreted in terms of increased WM integrity, FA may also increase in crossing fiber regions if one of the fiber population is degraded, and may be inferred based on the conjunction of MO changes is the same region^11^. Further, f1 and f2 models the evidence for two underlying fiber populations and the joint multivariate pattern revealed by LICA-09 thus seems sensitive to the underlying anatomy at the intersection of two major WM pathways in this region. The robust clinical and cognitive relevance of this component was supported by significant univariate associations with a wide range of cognitive and clinical domain scores (Fig. 2). Strikingly, associations indicated that the brain pattern in LICA-09 was less expressed in individuals with more severe general psychopathology, and more expressed in individuals with higher neuropsychological performance level. The reduction in subject weights for LICA-09 with increasing psychopathology implies a reduction of crossing fibers in the same area for individuals with increased symptom burden (Supplementary Fig. 5). Although LICA-09 showed a significant negative association with age, its associations with cognitive and clinical features were relatively stable across the sampled age-range. Importantly, Supplementary Fig. 6 illustrates that the general cognitive score and psychopathology exhibited independent associations with LICA-09. This WM microstructural pattern is of particular interest with respect to mental health for several reasons: UF is among the tracts with the highest heritability^33^, it displays a protracted maturation pattern as compared to other fiber tracts^8^, and is hypothesized to subserve a limbic temporo-amygdala-orbitofrontal network, critically involved in integration of emotional states with cognition and behavior^32, 34^. When investigating individual clinical ICA components, each of which express more symptom specificity compared to the general psychopathology score, similar relationships with LICA-09 subject weights (Supplementary Fig. 7) was observed for anxiety and harm avoidance (IC2), antisocial and norm violating behavior (IC3) as well as for psychosis prodrome and psychotic symptoms (IC4). This confirms that the reported brain pattern is broadly associated with a range of psychopathology symptoms rather than showing specificity with a certain clinical domain. These three symptom-categories have been shown to belong to separate empirical symptom-factors, often termed internalizing, externalizing and thought-disorder symptoms, however they also load on a general psychopathology factor implying shared common risk^3^. The present results point to altered limbic temporo-amygdala-orbitofrontal pathways as a candidate for such a transdiagnostic brain phenotype in psychiatric disease.

**Figure 2:**
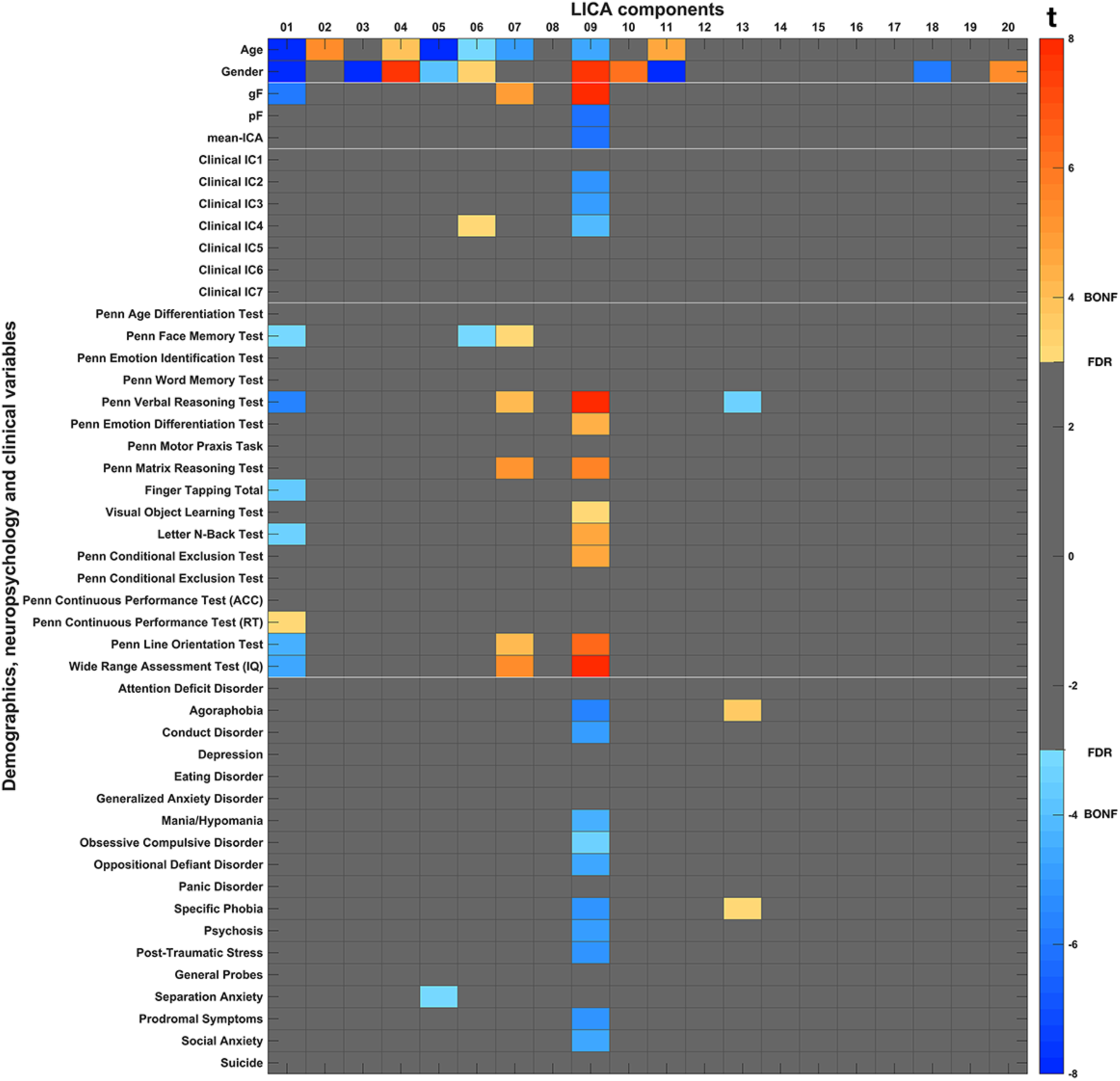
Univariate results for the subject weights of the 20 LICA components supports the cognitive and clinical relevance of LICA-09. All clinical and cognitive associations are adjusted for age, gender and tSNR, and corrected for multiple comparisons using a false discovery rate (FDR) of q=.05. The brain pattern captured by LICA-09 shows robust and inverse associations with cognitive and clinical scores, both for the gF/mean-ICA and ICA component scores, as well as for individual cognitive and clinical domain scores.

Fronto-temporal connectivity has been implicated in a range of mental illnesses including schizophrenia and its prodrome^35^, conduct disorder and psychopathy^31^ as well as anxiety and harm avoidance^36, 37^. The current findings show that such abnormalities are present also in non-diagnosed, but putatively at-risk youth. The distinct brain pattern predicting cognitive and clinical traits captures the developmental coordination of two major white matter tracts during late childhood and early adolescence, accompanying global age-related maturation of brain WM. The current work comes with some limitations. The size of the available sample for the heritability analysis in the PNC sample is on the lower end, and so point estimates should be interpreted with caution as the confidence intervals are wide. However, the estimated negative genetic correlation between cognition and psychopathology is unlikely to occur in a population with zero or positive values for rG, and the sign of the correlation is also in agreement with large-scale studies on cognition and psychopathology in adult populations^38^. <u>F</u>uture studies with larger samples are needed to determine whether there is also a genetic correlation between the observed brain phenotype and psychopathology. However, its robust associations with heritable general psychopathology and general cognitive factors warrant further investigation of fronto-temporal connectivity as an intermediate brain based phenotype in mental illness, and serve as a proof-of-principle for the emerging concept that a healthy transition from childhood to adolescence is dependent on an integrated yet partly independent development of distinct WM pathways in the brain.

## Acknowledgements

The authors were funded by the Research Council of Norway (#213837, #223273, #229129, #204966/F20, #249795, #251134), the South-Eastern Norway Regional Health Authority (#2017-112, #2014-097, #2015-073), KG Jebsen Stiftelsen, and the European Commission 7th Framework Programme (#602450, IMAGEMEND). Support for the collection of the data sets was provided by grant RC2MH089983 awarded to Raquel Gur and RC2MH089924 awarded to Hakon Hakonarson. All subjects were recruited through the Center for Applied Genomics at The Children’s Hospital in Philadelphia.

## Supplementary figures and tables

**Supplementary Figure 1.**
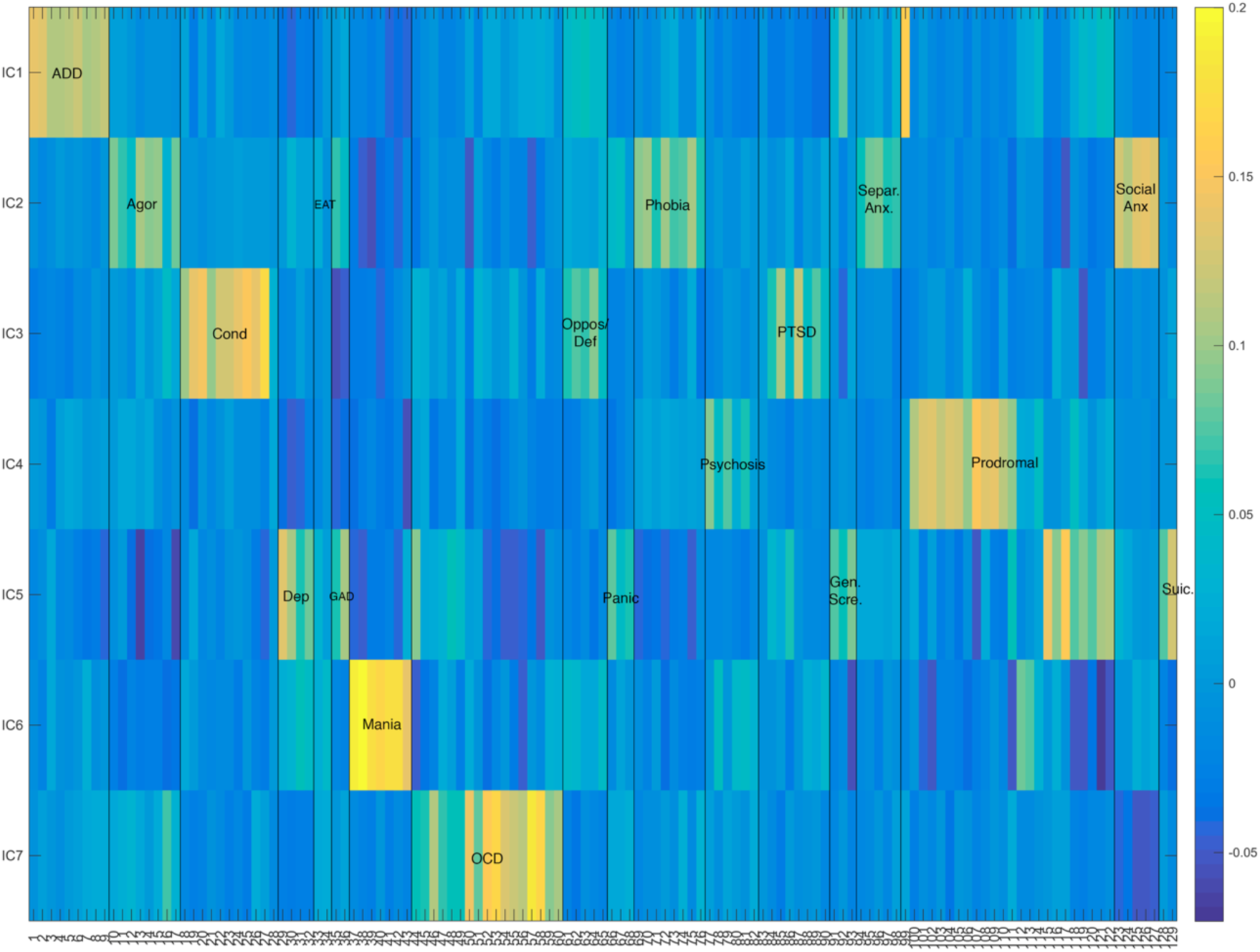
The weights for each of the 129 clinical questionnaire items on the seven estimated independent components. Clinical IC1 captures variance related to the attention deficit disorder (ADD) items, but also the attention item from the prodromal symptom severity scale (SIPS). IC2 encompasses items across several anxiety questionnaires, with highest weighting for social anxiety, but also agoraphobia, specific phobias and separation anxiety. IC3 is related to norm violating behaviours. Highest weights are assigned to conduct disorder items and oppositional defiance disorder, but also PTSD items related to experiences of violent behaviour. IC4 includes items from the SIPS questionnaires related to positive symptoms and delusions, as well as items on the psychosis questionnaire related to hallucinations. IC5 captures variance related to items across several clinical domains, including depression, generalized anxiety disorder, panic disorder, the negative symptoms of the SIPS questionnaire, as well as suicidal ideation. Component six (IC6) is related to mania and items capturing episodes of increased energy, reduced need for sleep, and increased speed of thought and speech. Component seven (IC7) is related to obsessive-compulsive disorder, and has the highest weights for items related to repetitive behaviours of counting, arranging and cleaning, but also to intrusive thoughts such worrying about germs/contamination. Color scale represents item loading on the independent components.

**Supplementary Figure 2:**
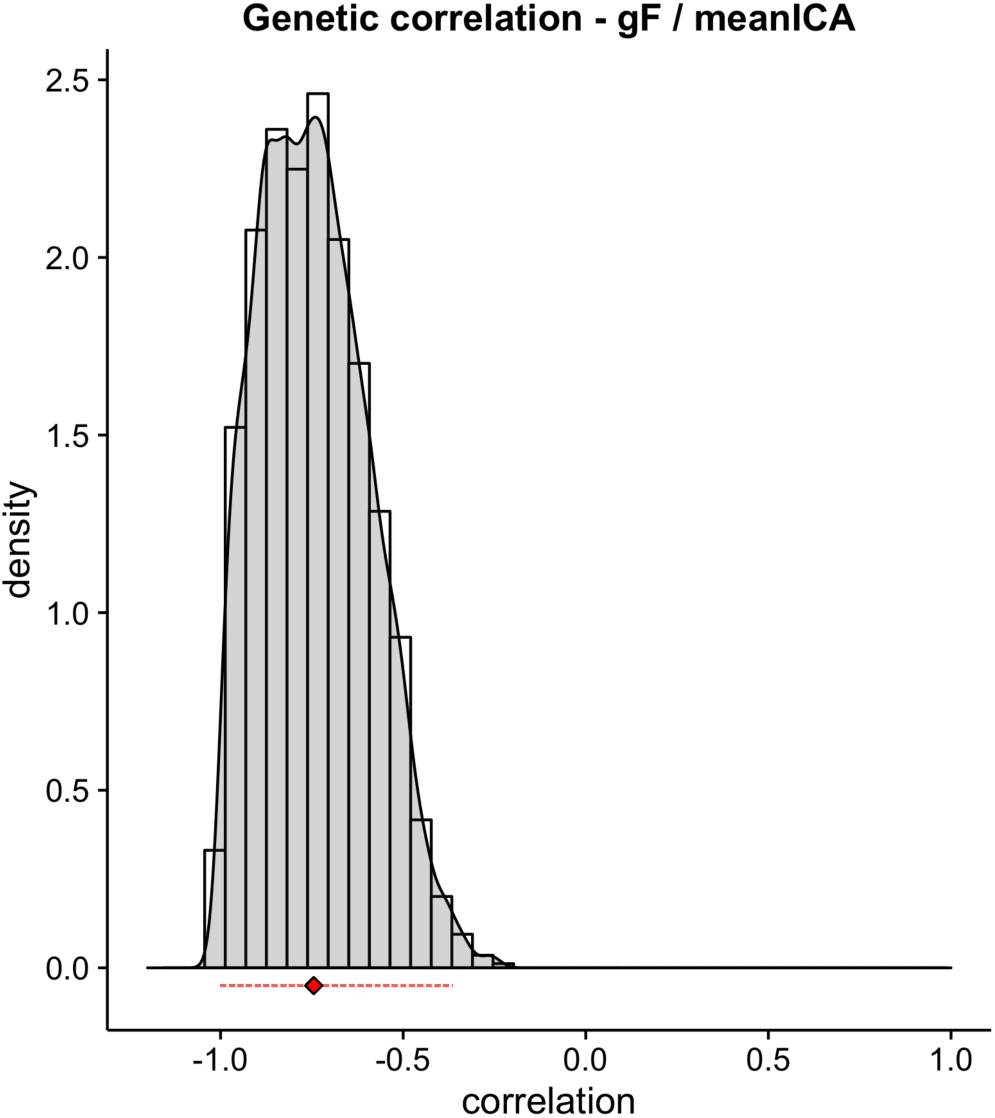
To get an estimate of the precision for the point estimate of the genetic correlation (rG) between gF and mean-ICA-scores, we employed resampling by drawing 6000 random subsamples (n=2,746) from the full sample (n=2,945), without replacement. The density plot shows the distribution of rG-estimates across sub-samples. The mean genetic correlation (rG) was r=−.74, s.e.=.15 and the 99% CI was [−1, −.36], indicated by the red diamond and line, respectively.

**Supplementary Figure 3:**
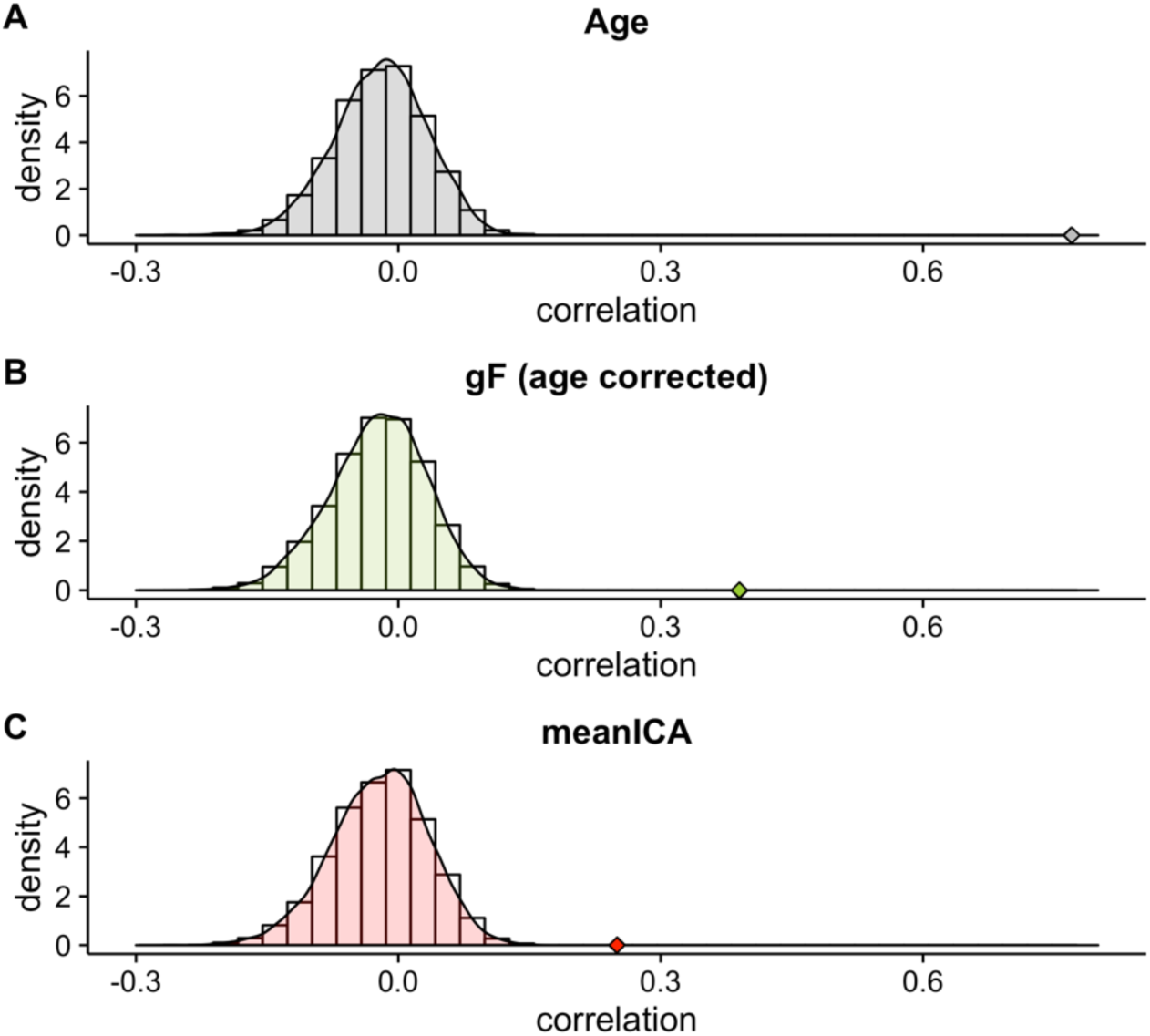
For each of the predicted traits (Age, gF and mean-ICA) we performed 10,000 permutations to assess whether the correlation between true and predicted scores where above chance level. Density plots represent the null-distributions, diamonds indicate the observed correlations. For Age and gF none of the permuted values exceeded the observed correlations, while the permuted p-value for mean-ICA prediction was .0001 (two-tailed).

**Supplementary Figure 4:**
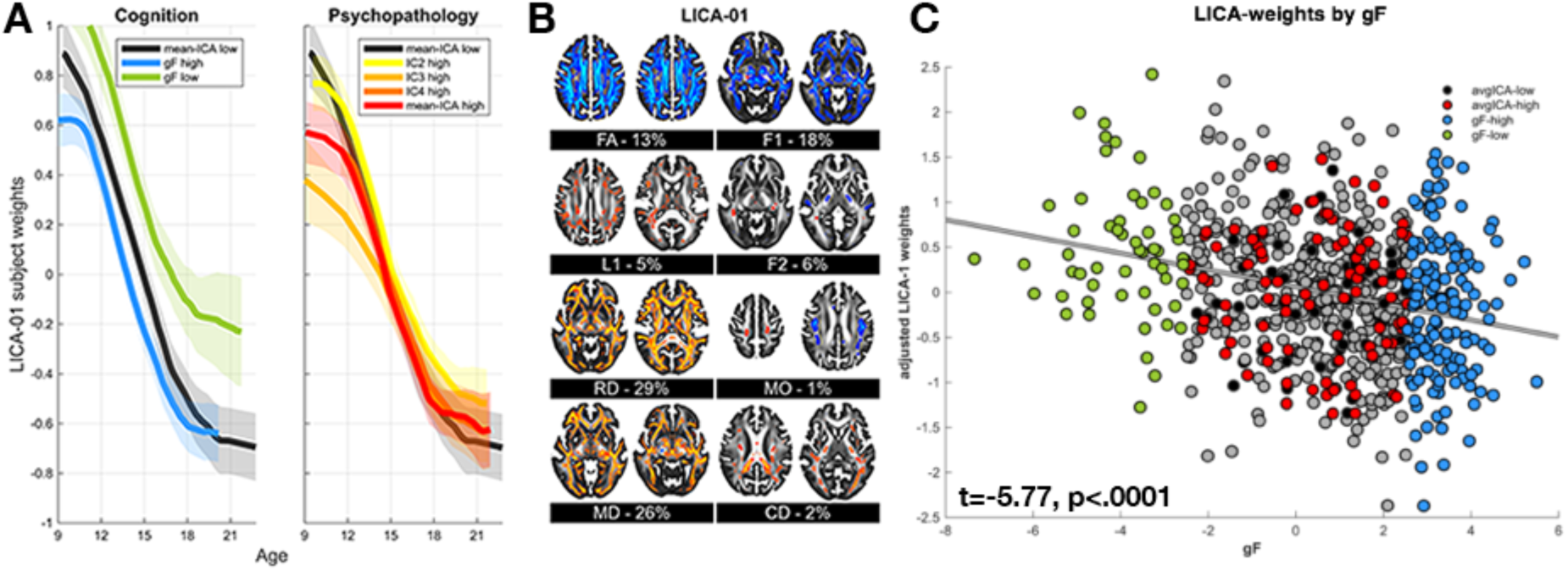
LICA-01 captures spatially global changes across the age-span. **(A)** Subject weights decrease as a function of age. **(B)** This indicates an increase predominantly in FA, f1, RD and MD (percentages represent the modalities contribution to the LICA-component) during neurodevelopment. Spatial map thresholded z=3. **(C)** A negative association between gF and LICA-01 subject weights indicate higher FA, f1, RD and MD in high performing individuals.

**Supplementary Figure 5:**
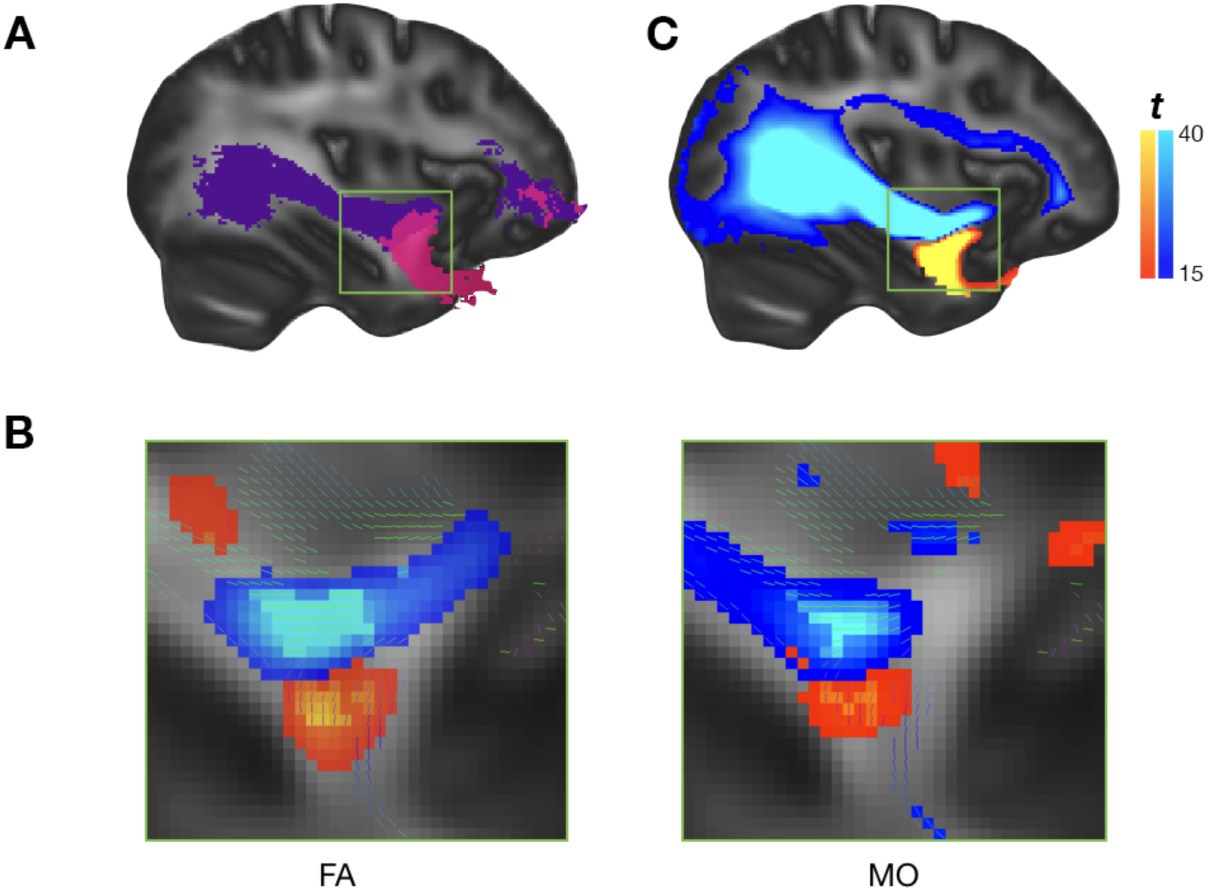
Seed based probabilistic tractography confirm involvement of UF. **(A)** To confirm that the crossing fiber area in the insular region is indeed related to intersection of IFOF (purple) and UF (pink) fibers (left panel, MNI x-coordinate=35 mm), we performed a seed-based probabilistic whole-brain tractography from ROIs based on LICA-09 weights in this region. **(B)** The seeds were based on voxels showing a conjunction of changes in both FA and MO (thresholded at z=3), an approach previously reported as sensitive to crossing fiber regions. The cluster with negative LICA spatial weights (blue) represent the insular UF-region with increased FA and MO with higher symptom burden while the more inferior cluster with positive values (red) represent a more inferior part of the UF showing the opposite pattern clinical pattern. **(C)** When contrasting their whole-brain connectivity probabilities the former cluster shows higher connectivity probability with the IFOF while the latter shows higher connectivity probability with the UF.

**Supplementary Figure 6:**
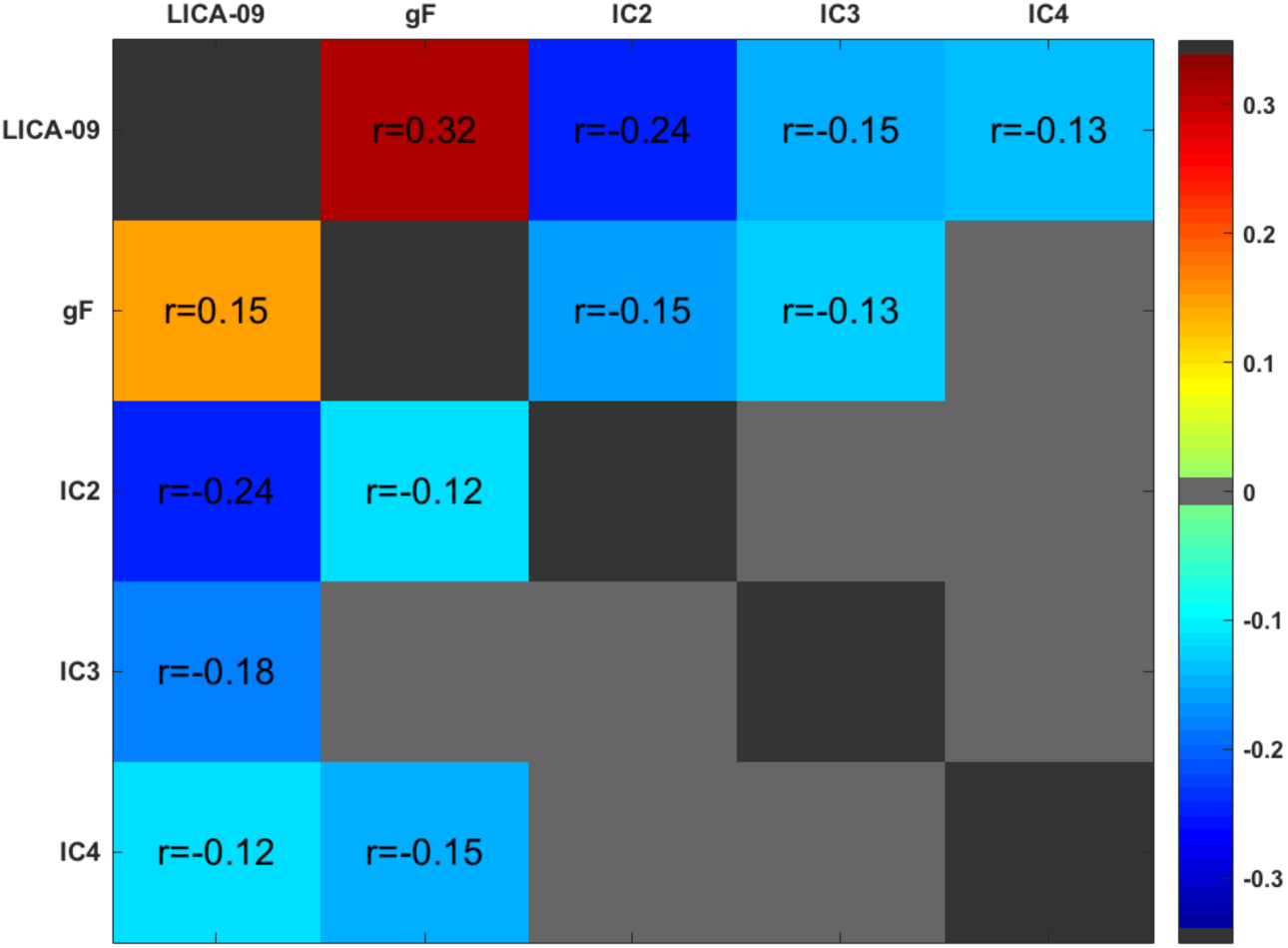
The full and partial correlations (including adjustment for age, gender and tSNR) between LICA-09 subject weights, gF, and clinical IC2 (anxiety and harm avoidance), IC3 (norm violating behaviour) and IC4 (psychosis prodrome) scores. Matrix threshold at q<.05 (FDR).

**Supplementary Figure 7:**
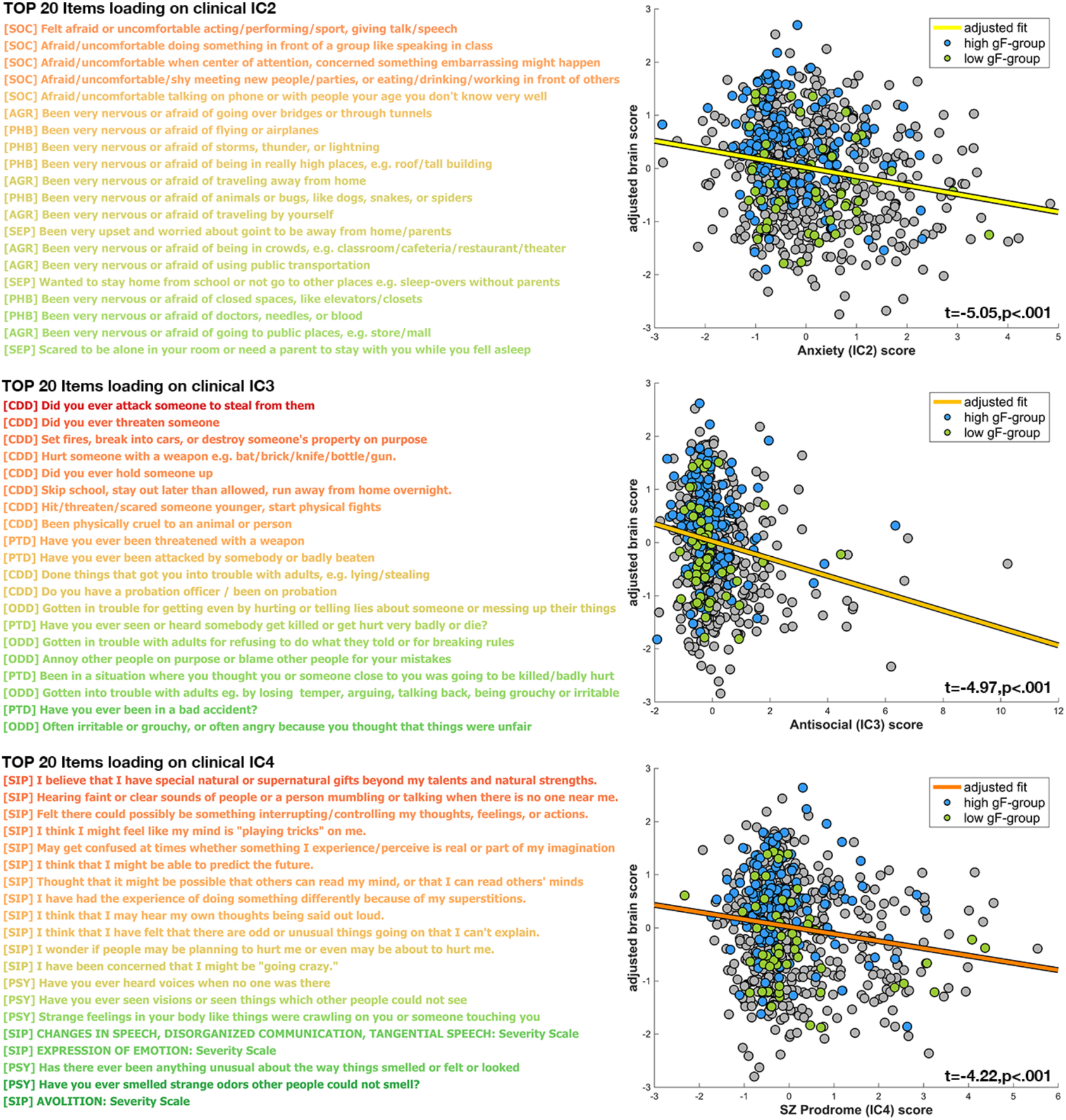
LICA-09 is associated with a wide range of clinical symptoms. Three of the clinical ICA-components showed an association with LICA-09 subject weights. Left panels shows the 20 highest loading clinical questionnaire items for clinical IC2 (anxiety and harm avoidance), IC3 (antisocial and norm violating behavior) and IC4 (psychosis prodrome). Warm colors represent higher weights. Right panel displays the LICA-09 subject weights regressed against the clinical ICA scores, adjusted for age, gender and tSNR.

**Supplementary Table 1:** General cognitive ability and general symptom burden are linked to common SNPs. SNPs explained 18% of the variability in gF scores and 16% of the variability in mean-ICA scores. V(G)/Vp is the estimated phenotypic variability attributable to additive effects of SNPs. r(g) is the estimated genetic correlation between phenotypes.

| SNP heritability |  |  |  |  |
| --- | --- | --- | --- | --- |
| Trait | N | V(G)/Vp | s.e. | p |
| gF (cognitive) | 2946 | 0.183 | 0.09 | 0.013 |
| mean-ICA (clinical) | 2946 | 0.156 | 0.095 | 0.049 |
| Genetic correlation |  |  |  |  |
| Traits | N | r(g) | se | p (one-tailed) |
| gF/mean-ICA | 5893 | -0.81 | 0.51 | 0.032 |

**Supplementary Table 2:**
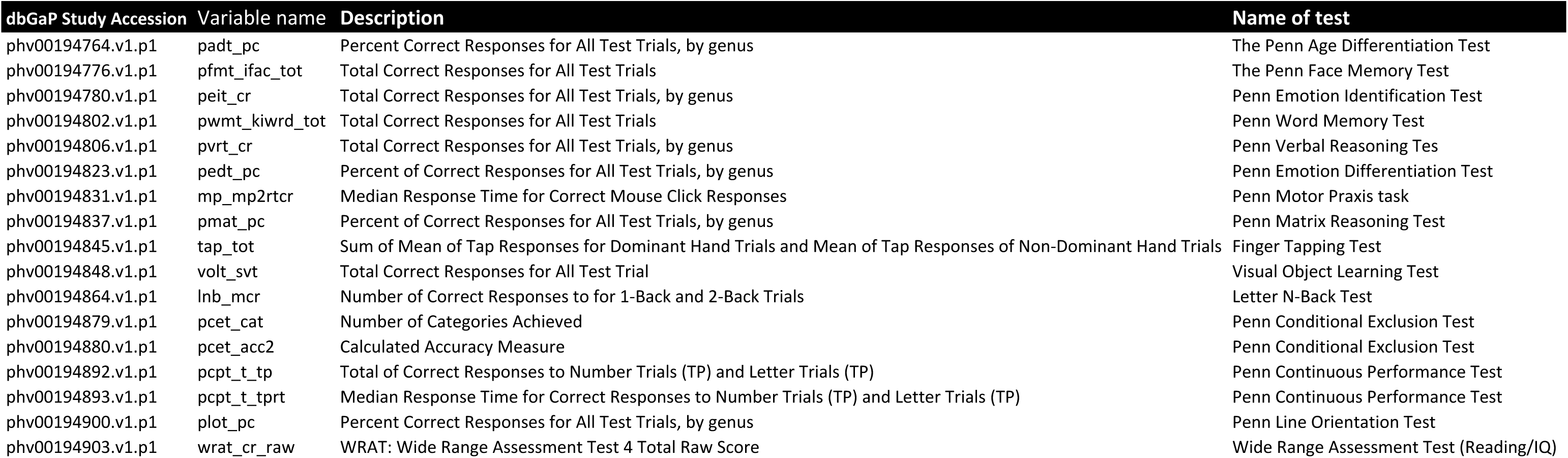
List of variables included in the PCA to compute the general cognition score (gF)

**Supplementary Figure 3:** List of the 129 clinical variables that entered the independent component analysis

| dbGaP Study Accession | Variable name | Item question |
| --- | --- | --- |
| phv00194007.v1.p1 | INT_TYPE | AP=Adult Proband (ages 18 and up); MP= Middle Proband (ages 11-17); MI=Middle Informant; YPI= Young Proband Informant (for ages 8-10) |
| **Attention Deficit Disorder (ADD)** |  |  |
| phv00194008.v1.p1 | ADD011 | Did you often have trouble paying attention or keeping your mind on your school, work, chores, or other activities that you were doing? |
| phv00194009.v1.p1 | ADD012 | Did you often have problems following instructions and often fail to finish school, work, or other things you meant to get done? |
| phv00194010.v1.p1 | ADD013 | Did you often dislike, avoid, or put off school or homework (or any other activity requiring concentration) |
| phv00194011.v1.p1 | ADD014 | Did you often lose things you needed for school or projects at home (assignments or books) or make careless mistakes in school work or other activities? |
| phv00194012.v1.p1 | ADD015 | Did you often have trouble making plans, doing things that had to be done in a certain kind of order, or that had a lot of different steps? |
| phv00194013.v1.p1 | ADD016 | Did you often have people tell you that you did not seem to be listening when they spoke to you or that you were daydreaming? |
| phv00194017.v1.p1 | ADD020 | Did you often have difficulty sitting still for more than a few minutes at a time, even after being asked to stay seated, or did you often fidget with your hands or feet or wiggle in your seat or were you "always on the go"? |
| phv00194018.v1.p1 | ADD021 | Did you often blurt out answers to other people's questions before they finished speaking or interrupt people abruptly? |
| phv00194019.v1.p1 | ADD022 | Did you often join other people's conversations or have trouble waiting your turn (e.g., waiting in line, waiting for a teacher to call on you in class)? |
| **Agoraphobia (AGR)** |  |  |
| phv00194035.v1.p1 | AGR001 | Have you ever been very nervous or afraid of: being in crowds (for example, a classroom, cafeteria, restaurant, or movie theater)? |
| phv00194036.v1.p1 | AGR002 | Have you ever been very nervous or afraid of: going to public places (such as a store or shopping mall)? |
| phv00194037.v1.p1 | AGR003 | Have you ever been very nervous or afraid of: being in an open field? |
| phv00194038.v1.p1 | AGR004 | Have you ever been very nervous or afraid of: going over bridges or through tunnels? |
| phv00194039.v1.p1 | AGR005 | Have you ever been very nervous or afraid of: traveling by yourself? |
| phv00194040.v1.p1 | AGR006 | Have you ever been very nervous or afraid of: traveling away from home? |
| phv00194041.v1.p1 | AGR007 | Have you ever been very nervous or afraid of: traveling in a car? |
| phv00194042.v1.p1 | AGR008 | Have you ever been very nervous or afraid of: using public transportation like a bus or SEPTA? |
| **Conduct disorder (CDD)** |  |  |
| phv00194062.v1.p1 | CDD001 | Was there ever a time when you often did things that got you into trouble with adults like lying or stealing (something worth more than \$5, from family, others, or stores)? |
| phv00194064.v1.p1 | CDD002 | Did you ever skip school, stay out at night later than you were supposed to (more than 2 hours), or run away from home overnight? |
| phv00194066.v1.p1 | CDD003 | Did you ever set fires, break into cars, or destroy someone else's property on purpose? |
| phv00194067.v1.p1 | CDD004 | Do you have a probation officer or have you ever been on probation? |
| phv00194068.v1.p1 | CDD005 | Did you often bully others (hitting, threatening or scaring someone who was younger or smaller), threaten or frighten someone on purpose, or often start physical fights with others? |
| phv00194069.v1.p1 | CDD006 | Have you ever been physically cruel to an animal or person (on purpose)? |
| phv00194070.v1.p1 | CDD007 | Did you ever: try to hurt someone with a weapon (a bat, brick, broken bottle, knife, or gun)? |
| phv00194071.v1.p1 | CDD008 | Did you ever: threaten someone? |
| phv00194072.v1.p1 | CDD009 | Conduct Disorder: Did you ever: hold someone up? |
| phv00194073.v1.p1 | CDD010 | Conduct Disorder: Did you ever: attack someone to steal from them? |
| phv00194074.v1.p1 | CDD011 | Did you ever: trick or threaten someone into having sex with you, or did anyone ever accuse you of making them do something sexual? |
| **Depression (DEP)** |  |  |
| phv00194091.v1.p1 | DEP001 | Has there ever been a time when you felt sad or depressed most of the time? |
| phv00194092.v1.p1 | DEP002 | Has there ever been a time when you cried a lot, or felt like crying? |
| phv00194094.v1.p1 | DEP004 | Has there ever been a time when you felt grouchy, irritable or in a bad mood most of the time; even little things would make you mad? |
| phv00194096.v1.p1 | DEP006 | Has there ever been a time when nothing was fun for you and you just weren't interested in anything? |
| **Eating disorder (EAT)** |  |  |
| phv00194133.v1.p1 | EAT001 | Was there ever a time when you felt really fat or heavy, but other people said that you were too thin? |
| phv00194139.v1.p1 | EAT007 | Has there been a time when your eating was out of control - you'd eat a large amount of food in a short period of time and could not stop yourself? |
| **Generalized anxiety (GAD)** |  |  |
| phv00194149.v1.p1 | GAD001 | Have you ever been a worrier? |
| phv00194150.v1.p1 | GAD002 | Did you worry a lot more than most children/people your age? |
| **Mania/Hypomania (MAN)** |  |  |
| phv00194185.v1.p1 | MAN001 | Have there been times when you were much more active, excited or energetic than usual, had problems sitting still, or needed to move around a lot? |
| phv00194186.v1.p1 | MAN002 | Has there ever been a time when you felt so full of energy that you couldn't stop doing things and didn't get tired? |
| phv00194187.v1.p1 | MAN003 | Has there ever been a time when you felt like you hardly needed sleep? |
| phv00194188.v1.p1 | MAN004 | Have there been times when you kept talking a lot, couldn't stop talking, talked faster than usual, had thoughts faster than usual, or had so many ideas in your head that you could hardly keep track of them? |
| phv00194189.v1.p1 | MAN005 | Have you ever had a time when you felt much more happy or excited than you usually do when there was nothing special going on? |
| phv00194190.v1.p1 | MAN006 | Have you ever had a time when you felt like you could do almost anything? |
| phv00194191.v1.p1 | MAN007 | Has there ever been a time when you felt unusually grouchy, cranky, or irritable; when the smallest things would make you really mad? |
| **Obsessive compulsive disorder (OCD)** |  |  |
| phv00194390.v1.p1 | OCD001 | Have you ever been bothered by thoughts that don't make sense to you, that come over and over again and won't go away, such as concern with harming others/self? |
| phv00194391.v1.p1 | OCD002 | Have you ever been bothered by thoughts that don't make sense to you, that come over and over again and won't go away, such as pictures of violent things? |
| phv00194392.v1.p1 | OCD003 | Have you ever been bothered by thoughts that don't make sense to you, that come over and over again and won't go away, such as thoughts about contamination/germs/illness? |
| phv00194393.v1.p1 | OCD004 | Have you ever been bothered by thoughts that don't make sense to you, that come over and over again and won't go away, such as fear that you would do something/say something bad without intending to? |
| phv00194394.v1.p1 | OCD005 | Have you ever been bothered by thoughts that don't make sense to you, that come over and over again and won't go away, such as feelings that bad things that happened were your fault? |
| phv00194395.v1.p1 | OCD006 | Have you ever been bothered by thoughts that don't make sense to you, that come over and over again and won't go away, such as forbidden/bad thoughts? |
| phv00194396.v1.p1 | OCD007 | Have you ever been bothered by thoughts that don't make sense to you, that come over and over again and won't go away, such as need for symmetry/exactness? |
| phv00194397.v1.p1 | OCD008 | Have you ever been bothered by thoughts that don't make sense to you, that come over and over again and won't go away, such as religious thoughts? |
| phv00194400.v1.p1 | OCD011 | Have you ever had to do something over and over again - that would have made you feel really nervous if you couldn't do it, like: cleaning or washing (for example, your hands, house)? |
| phv00194401.v1.p1 | OCD012 | Have you ever had to do something over and over again - that would have made you feel really nervous if you couldn't do it, like: counting? |
| phv00194402.v1.p1 | OCD013 | Have you ever had to do something over and over again - that would have made you feel really nervous if you couldn't do it, like: checking (for example, doors, locks, ovens)? |
| phv00194403.v1.p1 | OCD014 | Have you ever had to do something over and over again - that would have made you feel really nervous if you couldn't do it, like: getting dressed over and over again? |
| phv00194404.v1.p1 | OCD015 | Have you ever had to do something over and over again - that would have made you feel really nervous if you couldn't do it, like: going in and out a door over and over again? |
| phv00194405.v1.p1 | OCD016 | Have you ever had to do something over and over again - that would have made you feel really nervous if you couldn't do it, like: ordering or arranging things? |
| phv00194406.v1.p1 | OCD017 | Have you ever had to do something over and over again - that would have made you feel really nervous if you couldn't do it, like: doing things over and over again at bedtime, like arranging the pillows, sheets, or other things? |
| phv00194407.v1.p1 | OCD018 | Have you ever saved up so many things that people complained or they got in the way |
| phv00194408.v1.p1 | OCD019 | Do you feel the need to do things just right (like they have to be perfect)? |
| **Oppositional defiant disorder (ODD)** |  |  |
| phv00194431.v1.p1 | ODD001 | Was there a time when you often did things that got you into trouble with adults such as losing your temper, arguing with or talking back to adults, or being grouchy or irritable with them? |
| phv00194432.v1.p1 | ODD002 | Was there a time when you often got into trouble with adults for refusing to do what they told you to do or for breaking rules at home/school? |
| phv00194434.v1.p1 | ODD003 | Did you often annoy other people on purpose or blame other people for your mistakes (excluding siblings)? |
| phv00194436.v1.p1 | ODD005 | Did you ever get into trouble for getting even with other people by doing things to hurt them, telling lies about them, or messing up their things? |
| phv00194437.v1.p1 | ODD006 | Were you often irritable or grouchy, or did you often get angry because you thought that things were unfair? |
| **Panic disorder (PAN)** |  |  |
| phv00194448.v1.p1 | PAN001 | Have you ever had an attack like this? |
| phv00194449.v1.p1 | PAN003 | Has there ever been a time when all of a sudden you felt very, very scared or uncomfortable - and your chest hurt, you couldn't catch your breath, your heart beat very fast, you felt very shaky, and sweaty/tingly/numb in your hands or feet? |
| phv00194450.v1.p1 | PAN004 | Has there ever been a time when all of a sudden, you felt that you were losing control, something terrible was going to happen, that you were going crazy, or going to die? |
| **Specific phobia (PHB)** |  |  |
| phv00194468.v1.p1 | PHB001 | have you ever been very nervous or afraid of animals or bugs, like dogs, snakes, or spiders? |
| phv00194469.v1.p1 | PHB002 | have you ever been very nervous or afraid of being in really high places, like a roof or tall building? |
| phv00194470.v1.p1 | PHB003 | have you ever been very nervous or afraid of water or situations involving water, such as a swimming pool, lake, or ocean? |
| phv00194471.v1.p1 | PHB004 | have you ever been very nervous or afraid of storms, thunder, or lightning? |
| phv00194472.v1.p1 | PHB005 | have you ever been very nervous or afraid of doctors, needles, or blood? |
| phv00194473.v1.p1 | PHB006 | have you ever been very nervous or afraid of closed spaces, like elevators or closets? |
| phv00194474.v1.p1 | PHB007 | have you ever been very nervous or afraid of flying or airplanes? |
| phv00194475.v1.p1 | PHB008 | have you ever been very nervous or afraid of any other things or situations? |
| **Psychosis (PSY)** |  |  |
| phv00194490.v1.p1 | PSY001 | Have you ever heard voices when no one was there? |
| phv00194507.v1.p1 | PSY020 | Has there ever been anything unusual about the way things smelled or felt or looked? |
| phv00194518.v1.p1 | PSY029 | Have you ever seen visions or seen things which other people could not see? |
| phv00194531.v1.p1 | PSY050 | Have you ever smelled strange odors other people could not smell? |
| phv00194539.v1.p1 | PSY060 | Have you ever had strange feelings in your body like things were crawling on you or someone touching you and nothing or no one was there? |
| phv00194546.v1.p1 | PSY071 | Have you ever believed in things and later found out they weren't true, like people being out to get you, or talking about you behind your back, or controlling what you do or think? |
| **Post-traumatic stress (PTD)** |  |  |
| phv00194609.v1.p1 | PTD001 | Have you ever been in a flood or a tornado or an earthquake or a hurricane or some other natural disaster where you thought you were going to die or be seriously hurt? |
| phv00194610.v1.p1 | PTD002 | Have you ever been in a situation where you thought you or someone close to you was going to be killed or be hurt very badly (e.g. family violence)? |
| phv00194611.v1.p1 | PTD003 | Have you ever been attacked by somebody or badly beaten? |
| phv00194612.v1.p1 | PTD004 | Have you ever been very upset by someone forcing you to do something sexual? |
| phv00194614.v1.p1 | PTD006 | Have you ever been threatened with a weapon? |
| phv00194615.v1.p1 | PTD007 | Have you ever been in a bad accident? |
| phv00194616.v1.p1 | PTD008 | Other than television or at the movies, have you ever seen or heard somebody get killed or get hurt very badly or die? |
| phv00194617.v1.p1 | PTD009 | Have you ever been very upset by seeing a dead body or by seeing pictures of the dead body of somebody you knew well? |
| **General probes (SCR)** |  |  |
| phv00194639.v1.p1 | SCR001 | Have you ever talked to a counselor, psychologist, social worker, psychiatrist or some other professional about your feelings or problems with your mood or behaviors? |
| phv00194644.v1.p1 | SCR006 | Are you currently taking medication because of your emotions and/or behaviors? |
| phv00194645.v1.p1 | SCR007 | Have you ever had to go to a hospital and stay overnight because of problems with your mood, feelings, or how you were acting? |
| **Separation anxiety (SEP)** |  |  |
| phv00194654.v1.p1 | SEP500 | Since you were 5 years old, has there ever been a time when you had a lot of worries about your (attachment figures) and were very upset or got sick (for example, felt sick to your stomach, headaches, thrown-up) when you were away from him/her? |
| phv00194656.v1.p1 | SEP508 | Has there ever been a time when you wanted to stay home from school or not go to other places (for example, sleep-overs) without your (attachment figures)? |
| phv00194657.v1.p1 | SEP509 | When you knew that you were going to be away from home or (attachment figure(s)), did you get very upset and worry (e.g., when you learned (attachment figure(s)) were going on an upcoming trip or night out)? |
| phv00194658.v1.p1 | SEP510 | Did you ever worry/have bad dreams about something terrible happening to you or your (attachment figures) so that you would not see them again? |
| phv00194659.v1.p1 | SEP511 | Were you scared to be alone in your room (or any place in your house) or did you need your (attachment figure(s)) to stay with you while you fell asleep? |
| **Structural interview for prodromal symptoms (SIP)** |  |  |
| phv00194672.v1.p1 | SIP001 | TROUBLE WITH FOCUS AND ATTENTION Severity Scale |
| phv00194674.v1.p1 | SIP003 | I think that I have felt that there are odd or unusual things going on that I can't explain. |
| phv00194675.v1.p1 | SIP004 | I think that I might be able to predict the future. |
| phv00194676.v1.p1 | SIP005 | I may have felt that there could possibly be something interrupting or controlling my thoughts, feelings, or actions. |
| phv00194677.v1.p1 | SIP006 | I have had the experience of doing something differently because of my superstitions. |
| phv00194678.v1.p1 | SIP007 | I think I may get confused at times whether something I experience or perceive may be real or may be just part of my imagination or dreams. |
| phv00194679.v1.p1 | SIP008 | I have thought that it might be possible that other people can read my mind, or that I can read others' minds |
| phv00194680.v1.p1 | SIP009 | I wonder if people may be planning to hurt me or even may be about to hurt me. |
| phv00194681.v1.p1 | SIP010 | I believe that I have special natural or supernatural gifts beyond my talents and natural strengths. |
| phv00194682.v1.p1 | SIP011 | I think I might feel like my mind is "playing tricks" on me. |
| phv00194683.v1.p1 | SIP012 | I have had the experience of hearing faint or clear sounds of people or a person mumbling or talking when there is no one near me. |
| phv00194684.v1.p1 | SIP013 | I think that I may hear my own thoughts being said out loud. |
| phv00194685.v1.p1 | SIP014 | I have been concerned that I might be "going crazy." |
| phv00194698.v1.p1 | SIP027 | Do people ever tell you that they can't understand you? |
| phv00194699.v1.p1 | SIP028 | Do people ever seem to have difficulty understanding you? |
| phv00194700.v1.p1 | SIP030 | Changes in speech, disorganized communication, tangential speech Severity Scale |
| phv00194702.v1.p1 | SIP032 | Do you ever feel a loss of sense of self or feel disconnected from yourself or your life? |
| phv00194703.v1.p1 | SIP033 | Has anyone pointed out to you that you are less emotional or connected to people than you used to be? |
| phv00194704.v1.p1 | SIP035 | Changes in perception of self, others, or the world in general: Severity Scale |
| phv00194706.v1.p1 | SIP037 | EXPRESSION OF EMOTION: Severity Scale |
| phv00194707.v1.p1 | SIP038 | Within the past 6 months, are you having a harder time getting your work or schoolwork done? |
| phv00194708.v1.p1 | SIP039 | Within the past 6 months, are you having a harder time getting normal activities done? |
| phv00194709.v1.p1 | SIP041 | Occupational Functioning Severity Scale |
| phv00194711.v1.p1 | SIP043 | Avolition Severity Scale |
| **Social Anxiety (SOC)** |  |  |
| phv00194713.v1.p1 | SOC001 | was there ever a time in your life when you felt afraid or uncomfortable or really, really shy with people, like meeting new people, going to parties, or eating or drinking, writing or doing homework in front of others? |
| phv00194714.v1.p1 | SOC002 | was there ever a time in your life when you felt afraid or uncomfortable talking on the telephone or with people your own age who you don't know very well? |
| phv00194715.v1.p1 | SOC003 | was there ever a time in your life when you felt afraid or uncomfortable when you had to do something in front of a group of people, like speaking in class? |
| phv00194716.v1.p1 | SOC004 | was there ever a time in your life when you felt afraid or uncomfortable acting, performing, giving a talk/speech, playing a sport or doing a musical performance, or taking an important test or exam (even though you studied enough)? |
| phv00194717.v1.p1 | SOC005 | was there ever a time in your life when you felt afraid or uncomfortable because you were the center of attention and were concerned something embarrassing might happen and you felt very afraid or felt uncomfortable? |
| **Depression (DEP)** |  |  |
| phv00194734.v1.p1 | SUI001 | Have you ever thought a lot about death or dying? |
| phv00194735.v1.p1 | SUI002 | Have you ever thought about killing yourself? |

